# CGRig: a rigid-body protein model with residue-level interaction sites for long-time and large-scale protein assembly simulation

**DOI:** 10.64898/2026.03.21.713350

**Authors:** Yosuke Teshirogi, Tohru Terada

**Affiliations:** Department of Biotechnology, Graduate School of Agricultural and Life Sciences, The University of Tokyo, 1-1-1 Yayoi, Bunkyo-ku, Tokyo 113-8657, Japan

## Abstract

Molecular dynamics (MD) simulations are a powerful tool for investigating biomolecular dynamics underlying biological functions. However, the accessible spatiotemporal scales of conventional all-atom simulations remain limited by high computational costs. Coarse-graining reduces these costs by decreasing the number of interaction sites and enabling longer timesteps. In extreme cases, proteins are represented as single spherical particles; while such approximations facilitate cellular-scale simulations, they often sacrifice essential structural information, such as molecular shape and interaction anisotropy. Here, we present CGRig, a rigid-body protein model with residue-level interaction sites designed for long-time, large-scale simulations. In CGRig, each protein is treated as a single rigid-body embedding residue-level interaction sites. Its translational and rotational motions are described by the overdamped Langevin equation incorporating a shape-dependent friction matrix. Intermolecular interactions are calculated using Gō-like native contact potentials, Debye–Hückel electrostatics, and volume exclusion. We validated that CGRig accurately reproduces the translational and rotational diffusion coefficients expected from the friction matrix for an isolated protein. For dimeric systems, the model successfully maintained native complex structures. Furthermore, two initially separated proteins converged into the correct complex with an association rate consistent with all-atom simulations. Notably, CGRig achieved a simulation performance exceeding 17 μs/day for a 1,024-molecule system. These results demonstrate that CGRig provides an efficient framework for simulating protein assembly while retaining residue-level interaction specificity, making it a valuable tool for investigating large-scale biomolecular self-assembly.

## 1. INTRODUCTION

Molecular dynamics (MD) simulation is a powerful tool for elucidating the biomolecular dynamics that underlie biological functions^1^. By integrating the equations of motion with femtosecond time steps, MD simulations can capture microsecond-scale dynamics at atomistic resolution. Furthermore, they allow for the calculation of thermodynamic and kinetic quantities, providing access to physical properties that are often challenging to determine experimentally^2^. Driven by advances in hardware, such as GPUs^3^, and in simulation engines (e.g., GROMACS^4^, AMBER^5^, NAMD^6^, OpenMM^7^, LAMMPS^8^, and GENESIS^9^), MD simulation has become applicable to a wide range of biomolecules^10^.

Despite these successes, the spatiotemporal scales accessible to conventional all-atom (AA) MD simulations remain constrained by high computational costs^11^. The integration timestep is typically restricted to 2 fs, even when bonds involving hydrogen atoms are constrained^12,13^. Furthermore, the computational complexity grows with the number of atoms, severely limiting the size of the systems that can be modeled. Consequently, AA MD simulations are often practically limited to single proteins or small complexes within microsecond timescales^14^.

To extend the accessible spatiotemporal scales, two complementary strategies are commonly employed. The first is coarse-graining (CG), which reduces the number of interaction sites by mapping multiple atoms onto fewer effective particles^15,16^. A wide variety of CG protein models have been developed using diverse strategies, ranging from bottom-up^17–20^ and top-down approaches^21–26^ to recent machine-learning-based methodologies^27–31^. Martini is among the most widely used CG models, mapping approximately 2–4 non-hydrogen atoms to a single interaction site and thereby substantially reducing the number of interaction sites^32,33^. The second strategy involves extending the integration timestep by eliminating the fastest motions in the system. Because high-frequency intramolecular vibrations—particularly covalent-bond stretching—typically dictate the stability limits for explicit integration, constraining or removing these degrees of freedom can enable significantly larger timesteps.

A more radical reduction of internal degrees of freedom involves approximating entire proteins or domains as single spherical particles, whose motions are typically described by overdamped Langevin dynamics^34–37^. While such approximations drastically minimize the number of interaction sites and enable simulations to reach millisecond timescales or beyond—thereby facilitating studies of large-scale macromolecular self-assembly—they inevitably discard essential information regarding molecular shape and the anisotropic nature of protein–protein interactions. Furthermore, because protein association is intrinsically driven by the precise complementarity of surface geometries and residue-specific interactions, isotropic spherical models are inherently limited in their ability to provide mechanistic insights into molecular recognition and assembly^38,39^. Thus, bridging the gap between residue-level specificity and the need for massive spatiotemporal expansion remains a formidable challenge.

To overcome these limitations, we introduce CGRig, a novel coarse-grained rigid-body protein model. By treating each protein as a single rigid entity, CGRig eliminates high-frequency internal motions, allowing for significantly larger integration timesteps. Crucially, the model retains structural and chemical specificity by embedding residue-level interaction sites within the rigid frame, thereby capturing the anisotropic nature of protein–protein interactions. The translational and rotational dynamics are integrated using an overdamped Langevin formulation that incorporates a shape-dependent, full friction matrix, which enables translational–rotational coupling and frictional anisotropy for proteins of arbitrarily shape. Residue–residue interactions are parameterized in a bottom-up manner; native-contact interactions are optimized from AA reference simulations via force-matching, while non-native pairs are represented by excluded-volume repulsion. Additionally, long-range electrostatics are incorporated for charged residues. We validated the implementation and parameterization using an isolated ubiquitin and a set of protein dimer complexes. Finally, we demonstrate the applicability of CGRig to tubulin self-assembly by simulating a system containing 16 tubulin dimers.

## 2. THEORY

### 2.1. Numerical integration scheme

Long-time molecular motions are described by the overdamped Langevin equation. For a system of particles, the equation of motion for particle *i* is given by:

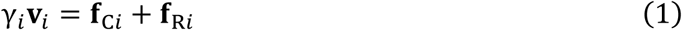

where *γ*_*i*_is the friction coefficient, **v**_*i*_is the velocity vector, and **f**_C*i*_and **f**_R*i*_denote the conservative and random forces, respectively. In a rigid-body system, each entity possesses both translational and rotational degrees of freedom. The equations of motion for rigid-body *i* in the laboratory frame are given by:

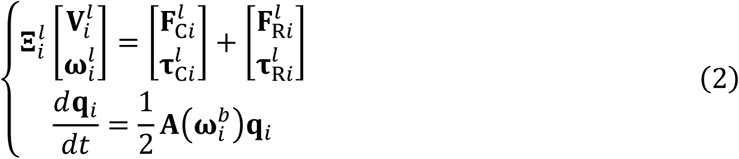

where 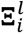 is the 6 × 6 friction matrix, 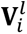 and 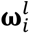 are the translational and angular velocities, 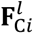 and 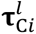 are the total force and torque vector resulting from conservative interactions, and 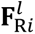 and 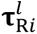 are the random force and torque vectors. The orientation of the body is described by the quaternion **q**_*i*_. Superscripts *l* and *b* denote laboratory and body-fixed frames, respectively. The matrix 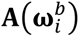 is defined using the angular velocity in the body-fixed frame:

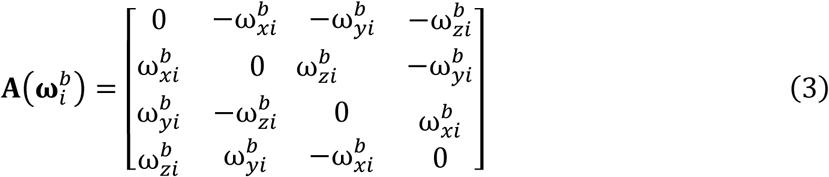

Since the friction matrix of a rigid-body is constant in the body-fixed frame, the equations of motion can be transformed as:

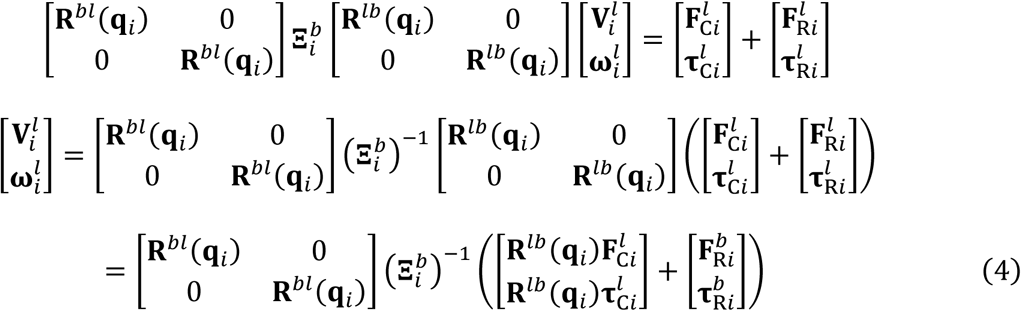

Here, **R**^*bl*^(**q**_*i*_) and **R**^*lb*^(**q**_*i*_) are 3 × 3 rotation matrices converting between the body-fixed and laboratory frames, with **R**^*lb*^(**q**_*i*_) = (**R**^*bl*^(**q**_*i*_))^−1^. The equations of motion are numerically integrated using an explicit scheme. The center-of-mass (COM) position **x**_*i*_and the orientation quaternion **q**_*i*_ of rigid-body *i* are updated as follows:

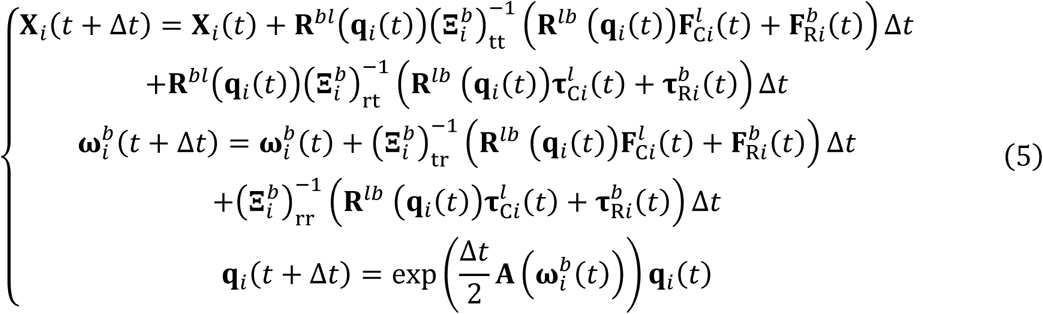

Here, the inverse of the friction matrix 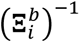 is partitioned into four 3 × 3 blocks, where the subscripts tt, rt, tr, and rr denote translation (top-left block), rotation-translation coupling (top-right block), translation-rotation coupling (bottom-left block), and rotation (bottom-right block), respectively. The random forces 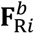 and the random torques 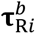 are generated from a normal distribution using the Cholesky factorization of the friction matrix:

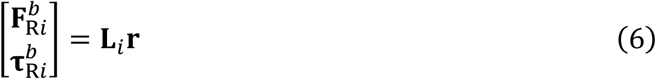

where **L**_*i*_ is the lower triangular matrix satisfying 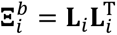. Each element of the 6-dimensional vector **r** follows a normal distribution with a mean of 0 and a standard deviation of 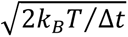, where *k*_*B*_is the Boltzmann constant and *T* is the absolute temperature. The friction matrix of each protein rigid-body is derived from the mobility matrix **D**_*i*_computed via US-SOMO^40,41^:

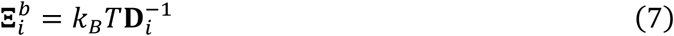

When a protein is approximated as an ellipsoidal or spherical body, the friction matrix 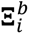 simplifies to a diagonal form^42^, and the rotation-translation coupling terms—represented by 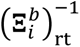 and 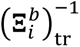 in Eq. (5)—vanish. For an ellipsoidal approximation, the diagonal elements satisfy 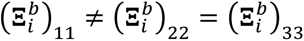 and 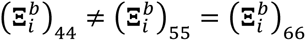, provided that the principal axis is aligned with the first coordinate. In the case of a spherical approximation, these elements further simplify to 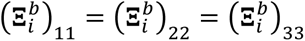 and 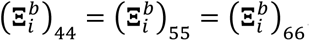.

### 2.2. Coarse-grained potential modeling

While the CGRig framework can accommodate interaction sites at various levels of resolution, we adopted a residue-level representation to balance physicochemical fidelity in protein–protein interactions with computational efficiency. In this model, an interaction site is placed at the Cα position of each residue. Although any residue-level coarse-grained interaction potential can be integrated into this framework, we developed an interaction potential—comprising the Go̅-like native contact potential, Debye–Hückel electrostatics, and volume exclusion—and compared its performance with established residue-level potentials. The total potential energy *U*(**x**, **q**) is defined as:

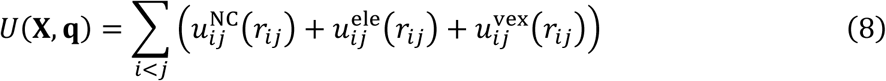

where 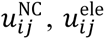 is the distance between interaction sites *i* and *j* . The terms 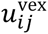, and represent the Go̅-like native contact, Debye–Hückel electrostatics, and volume exclusion potentials, respectively. We refer to this combined potential as the NELVEX (Native contact, Electrostatics, and Volume Exclusion) potential.

#### Native contact term

The native contact potential is defined based on a given reference structure as follows:

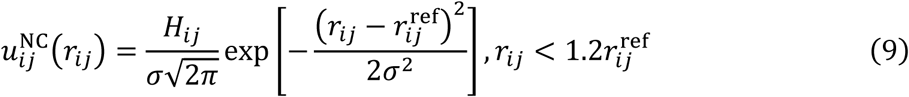

where 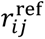 denotes the distance between interaction sites *i* and *j* in the reference structure. The parameter *H*_*ij*_determines the interaction strength, and the width parameter *σ* is uniformly set to 1.41421 Å. The coefficient *H*_*ij*_is assigned a non-zero value only for residue pairs with a Cα distance of less than 8 Å in the reference structure (i.e., native contact pairs); otherwise *H*_*ij*_ = 0 (non-native pairs). The matrix **H** is determined via the force-matching method^18,19^ by minimizing the following objective function:

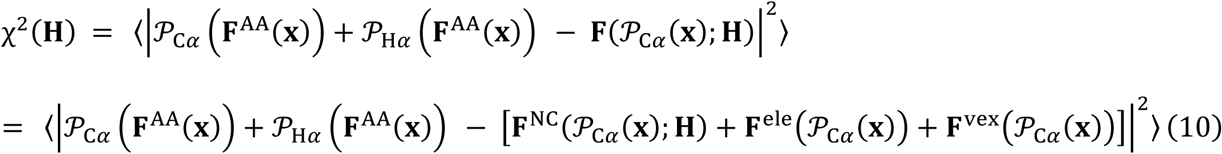

where **x** and **F**^AA^ represent the coordinates and the force vector of the AA model, respectively. The angular brackets 〈⋯ 〉 denote the ensemble average over the AA MD frames. The operators 𝒫_C*⍺*_(·) and 𝒫_H*⍺*_(·) extract the Cα and Hα atom components from an input vector, respectively. The term **F** = **F**^NC^ + **F**^ele^ + **F**^vex^ denotes the total force calculated from the potential terms in Eq. (8). Since the bond lengths between the Cα and Hα atoms are assumed to be constrained during the AA MD simulation, the force acting on an Hα atom is added to that of the corresponding Cα atom.

#### Electrostatics term

Electrostatic interactions between charged residues are described by the Debye–Hückel theory:

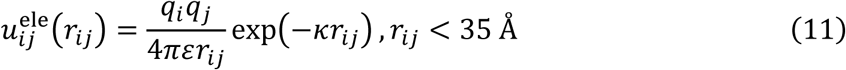

where *q*_*i*_ is the residue charge, *ɛ* is the dielectric constant of the solvent, and ***κ*** is the inverse Debye length. The potential is truncated at a cutoff distance of 35 Å.

#### Volume exclusion term

Volume exclusion interactions are calculated for all non-native pairs using a cosine-based potential:

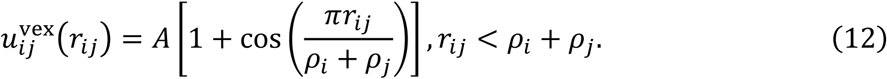

where *ρ*_*i*_ and *ρ*_*j*_ are the effective radii of residues *i* and *j*, respectively, which are taken from a previous study^43^. The parameter *A* is set to 2 kcal mol^−1^ for all pairs.

## 3. COMPUTATIONAL DETAILS

### 3.1. Simulation of an isolated protein system

To validate the numerical integration scheme of CGRig, we first applied the model to an isolated protein system. Ubiquitin (PDB ID: 1UBQ) was selected as the test case. The mobility matrix **D**_*i*_was calculated using US-SOMO^40,41^ and subsequently converted into the friction matrix 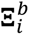 according to Eq. (7) . To evaluate the diffusion properties, we performed 100 independent replicas of rigid-body overdamped Langevin dynamics simulations with Δ*t* = 0.1 ps, each consisting of 10^6^ steps. These simulations were conducted using three different representations of the friction matrix: the full friction matrix, an ellipsoidal approximation, and a spherical approximation. In the ellipsoidal approximation, the two diagonal elements with similar values were averaged and used for the second and third axes in both the translational and rotational blocks. For the spherical approximation, the average of all three diagonal elements was used for each block. The translational and rotational diffusion coefficients were then calculated from the resulting trajectories to ensure consistency with the theoretical values derived from the friction matrix.

### 3.2. Simulations of protein dimer complexes Reference AA Simulations

To evaluate the stability of native complex structures across different CG potentials, we selected eleven protein dimer systems based on a previous study^44^: TβR-1–FKBP12 (PDB ID: 1B6C), barnase–barstar (1BRS), BPTI–MT-SP1 (1EAW), ImGP synthase (1GPW), CD2–CD58 (1QA9), ubiquitin–UEV domain (1S1Q), PEX5P–MSCP2 (2C0L), Ubq–Ubq-ligase (2OOB), BPTI–trypsin (2PTC), SGPB–OMTKY3 (3SGB), and colE7–Im7 (7CEI). Each complex was solvated in a cubic water box using the Solution Builder^45,46^ of the CHARMM-GUI server^47^. During this step, missing residues in the barnase–barstar (1BRS), ImGP synthase (1GPW), and PEX5P–MSCP2 (2C0L) systems were modeled using the server’s built-in tools. Potassium and chloride ions were added to a concentration of 0.15 M, except for the barnase–barstar system, which was set to 0.05 M. We employed the Amber ff19SB force field^48^ for proteins and the OPC water model^49^ for the solvent.

After energy minimization, each system was equilibrated in the *NVT* ensemble for 125 ps with a 1-fs timestep, followed by *NPT* equilibration for 100 ns with a 2-fs timestep. During the *NVT* equilibration, positional restraints were applied to protein non-hydrogen atoms with force constants of 400 and 40 kJ mol^−1^ nm^−2^ for backbone and side-chain atoms, respectively. The *NPT* equilibration was performed without any restraints. Subsequently, production simulations were carried out in the *NVT* ensemble for 500 ns, with snapshot recorded every 10 ps.

In all AA MD simulations, electrostatic interactions were calculated using the particle mesh Ewald (PME) method^50,51^. Van der Waals interactions were truncated at a cutoff distance of 9 Å. Bond lengths involving hydrogen atoms were constrained using the LINCS^13,52^ algorithm. The temperature was maintained at 303.15 K using a velocity-rescaling thermostat^53^, except for the barnase–barstar system, which was simulated at 298.15 K. During *NPT* equilibration, the pressure was maintained isotropically at 1×10^5^ Pa using a C-rescale barostat^54^. All AA MD simulations were performed using GROMACS 2023.2^4^.

#### CGRig Model Parameterization and Simulation

Reference structures for the CGRig model were determined by selecting the representative structure from the most populated cluster obtained through hierarchical clustering^55^ of the AA MD snapshots. The friction matrix 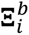 for each system was then calculated from these reference structures using US-SOMO^40,41^. The interaction coefficients **H** for the NELVEX potential were optimized by minimizing the objective function in Eq. (10) using the AA MD trajectories.

To benchmark the performance of the NELVEX potential, we performed five independent replicas of rigid-body overdamped Langevin dynamics simulations (10^6^ steps, Δ*t* = 0.01 ps) for each system. For comparison, identical simulations were conducted using established coarse-grained potentials: HPS-Urry^56^, KH^57^, and Mpipi^25^. Additionally, for the barnase–barstar system, protein–protein association simulations were initiated from a separated state (minimum distance > 35 Å). CGRig was implemented as an in-house plugin for LAMMPS 22Jul2025^8^, and all simulations were performed using this framework.

### 3.3. Tubulin assembly simulations and performance benchmarks

To evaluate the applicability of CGRig to large-scale biomolecular self-assembly, we applied the model to a microtubule-forming tubulin system. For parameterization, we utilized the structure of a 12-mer tubulin complex (comprising six α-tubulin and six β-tubulin subunits; PDB ID: 6EVW) (Figure S1A). In this structure, the α,β-bridging methylene carbon atom of GMPCPP was replaced with an oxygen atom to represent GTP. Missing residues were modeled using MODELLER^58^, and the C-termini at each subunit were capped with *N*-methyl groups. The force field parameters for GTP in the AA MD simulations were determined as follows: the molecular geometry was optimized, electrostatic potentials were calculated at the HF/6-31G(d) level of theory using Gaussian 16, revision C.02^59^. Partial charges were subsequently assigned using the restrained electrostatic potential (RESP) method^60^. Other force field parameters were taken from the general AMBER force field 2 (GAFF2)^61^. AA MD simulations were performed for this system under the same protocols and conditions as described in the previous section for the protein dimer complexes, with the exception of the production run length, which was set to 100 ns. In the CGRig framework, each α- and β-tubulin subunit was treated as a separate rigid-body, with residue-level interaction sites embedded at the Cα positions. Additionally, GTP molecules at the intermolecular interfaces were represented by five interactions sites, and Mg^2+^ ions were modeled as single interaction sites (Figure S1B). These additional sites were embedded within the corresponding protein rigid frame. The interaction coefficients **H** of the NELVEX potential were then determined for both longitudinal and lateral interfaces by optimizing the target function of Eq. (10) using the AA MD trajectory (see Figure S1A).

The resulting CGRig model was subsequently employed in assembly simulations of 16 tubulin dimers, which were initially arranged on a lattice. In addition to observing the assembly process, we conducted performance benchmarks to assess the computational efficiency and scalability of CGRig for large-scale systems. In these benchmarks, the number of tubulin dimers was varied, and computational acceleration was achieved by enabling CPU multithreading and GPU offloading via the OPENMP and KOKKOS plugins^62^, respectively. The benchmark calculations were carried out on a single node of the Miyabi supercomputer equipped with an NVIDIA Grace CPU and an NVIDIA H100 GPU.

## 4. RESULTS AND DISCUSSION

### 4.1. Validation of the CGRig framework using an isolated protein system

Figure 1 illustrates the workflow of the CGRig framework proposed in this study. To verify the numerical integration scheme, we first examined whether CGRig accurately reproduces the translational and rotational diffusion properties expected from the friction matrix of an isolated protein. We selected ubiquitin as a test system and calculated its full (6 × 6) friction matrix using US-SOMO^40,41^ based on the coordinates of its crystal structure. For comparison, friction matrices under ellipsoidal and spherical approximations were also derived.

**Figure 1.**
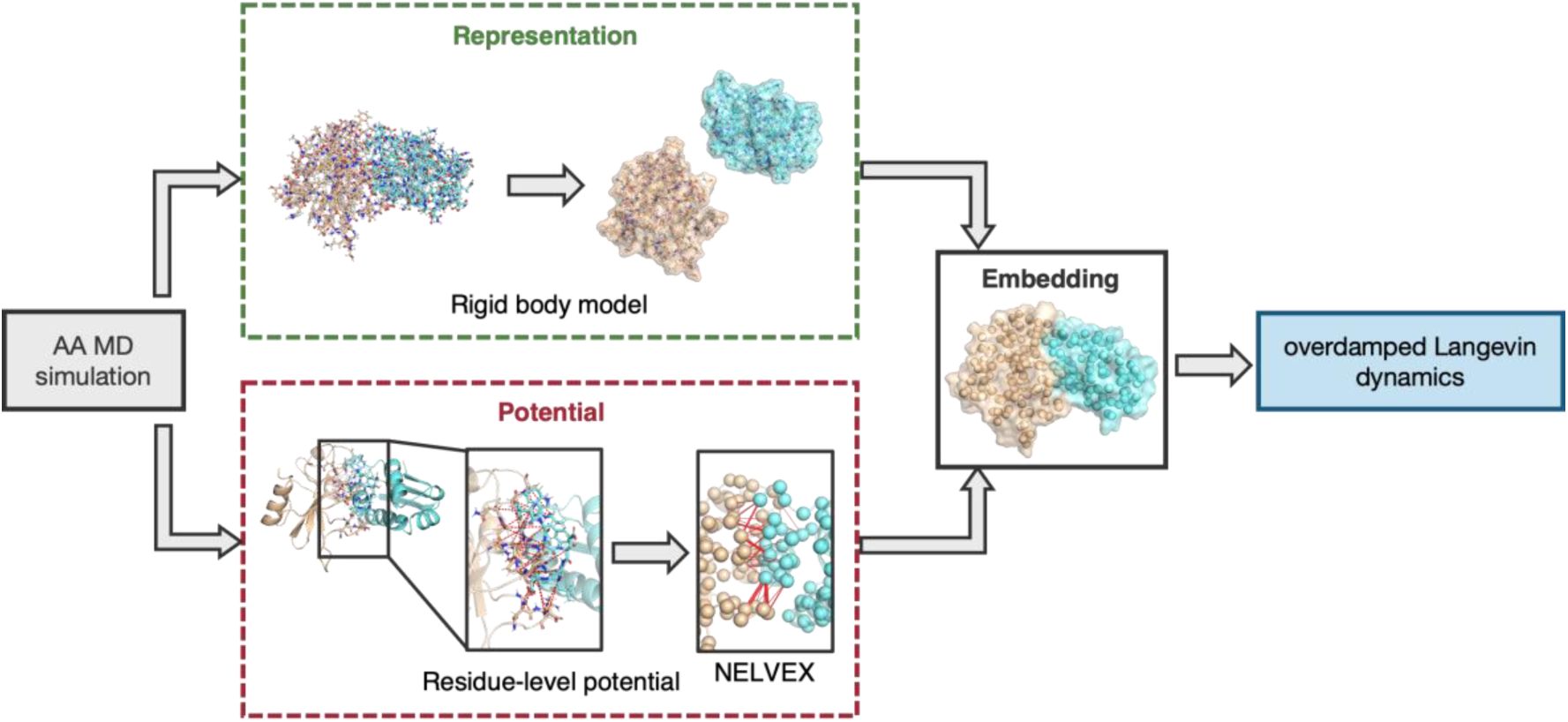
Overview of the CGRig framework. The workflow is illustrated using the barnase–barstar system. First, an all-atom (AA) MD simulation is conducted for the target complex. (Upper) Construction of a rigid-body representation from a representative structure of the AA MD simulation. (Lower) Parameterization of intermolecular interactions using the AA MD trajectory. In the magnified view, native-contact residue pairs are highlighted; in the final residue-level model, interaction sites at Cα positions are represented as spheres, with the thickness of the connecting lines reflecting the optimized interaction strengths (*H*_*ij*_). The rigid-body representation and the residue-level interaction model are combined by embedding the interaction sites within each rigid frame. The dynamics of the system are integrated using overdamped Langevin equations. All structural images were generated by PyMOL^63^.

Table 1 summarizes the translational (*D*_trans_) and rotational (*D*_rot_) diffusion coefficients along with the anisotropy, obtained from the CGRig simulations using the three different friction matrix representations. The *D*_trans_values calculated by CGRig were consistent across all representations (1.54–1.55 × 10^−6^ cm^2^ s^−1^) and showed excellent agreement with both the US-SOMO theoretical value (1.54 × 10^−6^ cm^2^ s^−1^) and experimental data (1.33–1.62 × 10^−6^ cm^2^ s^−1^)^64,65^. In contrast, the rotational diffusion properties were more sensitive to the friction matrix representation. The overall diffusion coefficients (*D*_rot_) and anisotropy values for the full friction matrix (3.95 × 10^7^ rad^2^ s^−1^ and 1.36) and the ellipsoidal approximation (3.94 × 10^7^ rad^2^ s^−1^ and 1.38) closely agreed with the US-SOMO theoretical values (3.94 × 10^7^ rad^2^ s^−1^ and 1.39), whereas the values for the spherical approximation showed significant deviations (3.83 × 10^7^ rad^2^ s^−1^ and 1.05). Furthermore, all the *D*_rot_ components (*D*_rot,1_, *D*_rot,2_, and *D*_rot,3_) for the full friction matrix closely matched those of US-SOMO. In the ellipsoidal approximation, however, *D*_rot,2_and *D*_rot,3_resulted in nearly identical values, failing to capture the difference between these components predicted by US-SOMO. This is because the ellipsoidal model by definition averages the corresponding diagonal elements of the friction matrix. These results demonstrate that the full friction matrix in the CGRig framework is essential for accurately describing the anisotropic diffusion behavior of proteins by consistently accounting for translation-rotation coupling.

**Table 1.** Calculated diffusion coefficients of isolated ubiquitin using different friction models^a^.

| Method | $D_{\text{trans}}$<br>($10^{-6}$<br>$\text{cm}^2 \text{s}^{-1}$) | $D_{\text{rot}}$ ($10^7 \text{ rad}^2$<br>$\text{s}^{-1}$)<sup>b</sup> | $D_{\text{rot},1}$<br>($10^7 \text{ rad}^2$<br>$\text{s}^{-1}$) | $D_{\text{rot},2}$<br>($10^7 \text{ rad}^2$<br>$\text{s}^{-1}$) | $D_{\text{rot},3}$<br>($10^7 \text{ rad}^2$<br>$\text{s}^{-1}$) | Anisotropy<sup>c</sup> |
| --- | --- | --- | --- | --- | --- | --- |
| CGRig (full) | $1.55 \pm 0.03$ | $3.95 \pm 0.10$ | $4.79 \pm 0.19$ | $3.61 \pm 0.16$ | $3.43 \pm 0.14$ | $1.36 \pm 0.06$ |
| CGRig<br>(ellipsoid) | $1.54 \pm 0.03$ | $3.94 \pm 0.09$ | $4.81 \pm 0.21$ | $3.51 \pm 0.14$ | $3.49 \pm 0.12$ | $1.38 \pm 0.07$ |
| CGRig (sphere) | $1.54 \pm 0.04$ | $3.83 \pm 0.08$ | $3.86 \pm 0.16$ | $3.79 \pm 0.15$ | $3.84 \pm 0.13$ | $1.05 \pm 0.03$ |
| US-SOMO<sup>40,41</sup> | 1.54 | 3.94 | 4.84 | 3.58 | 3.41 | 1.39 |
| Experiment<sup>64-69</sup> | 1.33 – 1.62 | 4.01 – 4.14 | – | – | – | 1.15 – 1.18 |

<sup>a</sup>Values for CGRig are presented as mean ± standard deviation
<sup>b</sup> $D_{\text{rot}} = (D_{\text{rot},1} + D_{\text{rot},2} + D_{\text{rot},3})/3$
<sup>c</sup>Anisotropy = $2D_{\text{rot},1}/(D_{\text{rot},2} + D_{\text{rot},3})$

### 4.2. Stability of protein dimer complexes

Next, we evaluated the stability of native complex structures in CGRig simulations by comparing the performance of our proposed NELVEX potential with established CG potentials, namely HPS-Urry^56^, KH^57^, and Mpipi^25^. Figure 2 displays the eleven protein dimer complexes examined in this study, along with the time evolutions of the Cα-RMSDs during the CGRig simulations using the different CG potentials. The RMSD plots reveal that the protein dimers rapidly dissociated and remained in an unbound state throughout the simulations when using the HPS-Urry, KH, and Mpipi potentials. In contrast, all complex structures remained stable with the NELVEX potential, exhibiting high consistency with the reference AA MD simulations (Figure S2). Although the 1QA9 complex exhibited larger RMSD values than the other complexes in the NELVEX simulations, this can be attributed to its relatively small binding interface (Figure S3), which allows for significant rotational fluctuation of one subunit against the other. Such fluctuations were also consistently observed in the reference AA MD simulations (Figure S2), suggesting that the increased RMSD reflects the inherent flexibility of the complex rather than an instability of the model. The existing CG potentials are designed for general-purpose applications and are parameterized based on the twenty standard amino acids. This implies that identical parameters are applied to the same residue types regardless of their structural positions. While these models have successfully described interactions in intrinsically disordered regions—such as in liquid–liquid phase separation phenomena—they often lack the specificity required to stabilize the complex structures of folded proteins. Therefore, the NELVEX potential, which incorporates structural-specific native contact terms, is highly recommended when applying CGRig to simulations of protein self-assembly phenomena.

**Figure 2.**
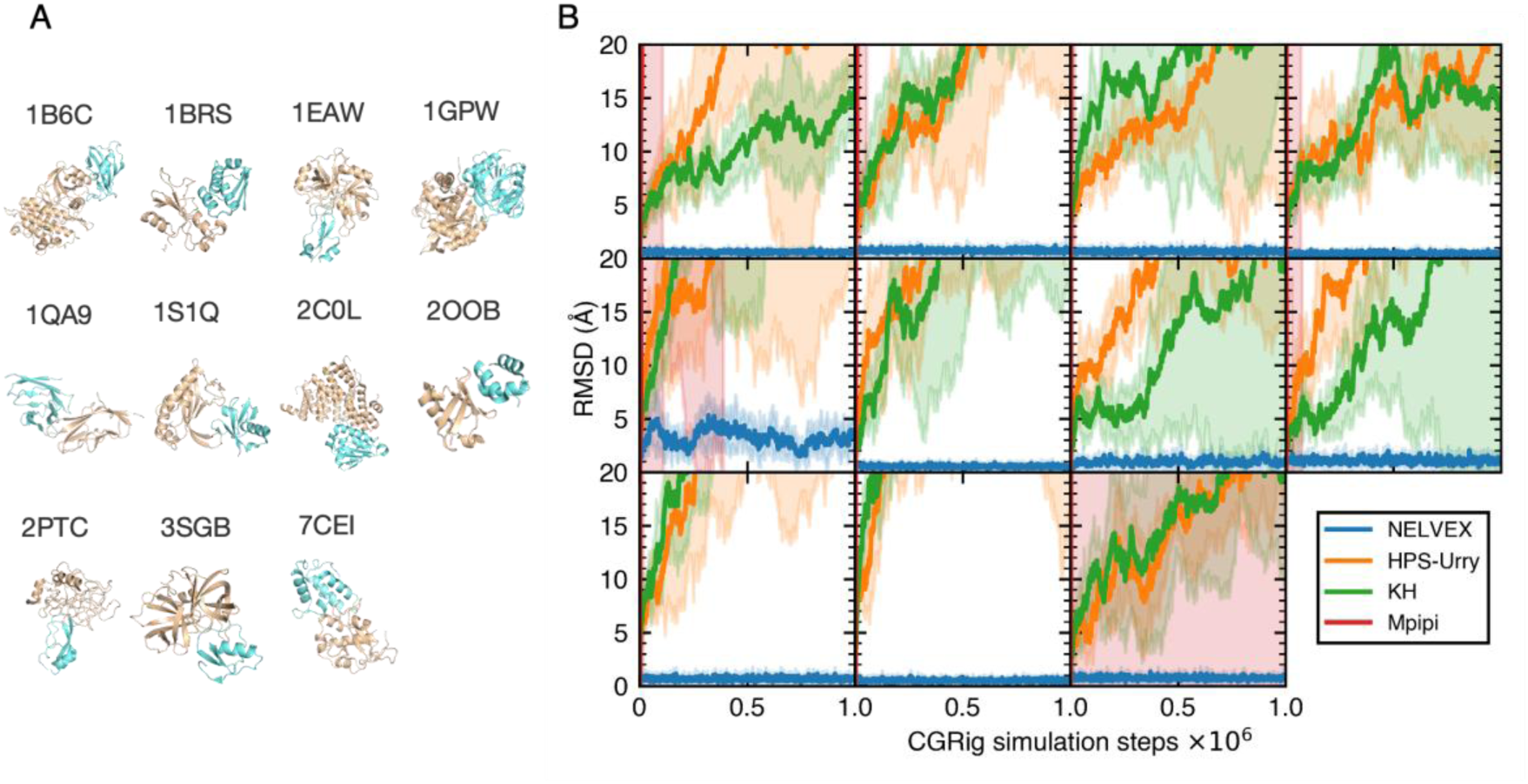
Validation of protein dimer simulations. (A) Structures of the tested complexes. For each complex, the larger subunit (longer amino acid sequence) is colored brown, and the smaller subunit (shorter sequence) is colored blue. (B) Time evolutions of Cα-RMSD for the smaller subunits, calculated after aligning the larger subunits to their respective reference structures. Solid lines represent the mean values of five independent runs, with the shaded areas indicating the standard deviation.

The interaction coefficients *H*_*ij*_can take either positive or negative values depending on the forces observed in the reference AA MD simulations, corresponding to repulsive and attractive interactions, respectively, when the distance *r*_*ij*_is greater than the reference distance *r*^ref^. This feature provides a clear contrast to conventional Gō-like potentials, in which only attractive forces are applied and a uniform coefficient is typically assigned to all native contact pairs. To examine whether the force-matched *H*_*ij*_parameters in the NELVEX formalism provides a clear advantage in terms of complex stability, we conducted five independent CGRig simulations (10^6^ steps, Δ*t* = 0.01 ps) for the barnase–barstar system using a single, uniform interaction strength (*H*_*ij*_ = *H*_const_) for all native contact pairs. The value of *H*_const_ was varied to investigate its effect on stability. Figure 3 compares the probability distributions of the Cα-RMSD obtained from the simulations using the NELVEX and conventional Gō-like potentials. In the simulations with the conventional Gō-like potentials, the probability of maintaining a near-native structure (Cα-RMSD < 1 Å) increased as the interaction strength was strengthened. However, the maximum probability achieved with the conventional Gō-like potential (0.793) remained lower than that achieved with the NELVEX potential (0.844). Moreover, when the interaction coefficients of the NELVEX potential were restricted to negative values (i.e., only attractive interactions), the complex stability was significantly degraded in 7 of the 11 systems (Figure S4). In particular, the proteins in the 1QA9 system underwent complete dissociation. These results indicate that both attractive and repulsive interactions between native contact pairs are essential for the stability of the complex. The force-matching scheme employed in this study automatically balances these forces to achieve optimal interactions. Based on these findings, the NELVEX potential was employed in all subsequent CGRig simulations throughout this study.

**Figure 3.**
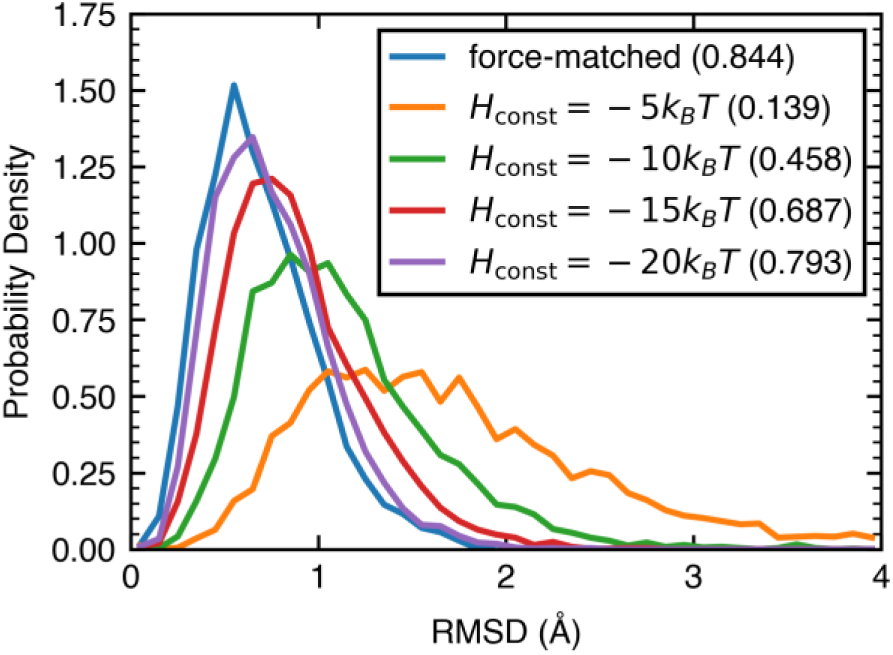
Comparison of Cα-RMSD probability distributions. The distributions were obtained from five independent CGRig simulations (10^6^ steps, Δ*t* = 0.01 ps) of the barnase–barstar system, comparing the NELVEX potential (force-matched *H*_*ij*_) with four models using different uniform, constant interaction strengths (*H*_*ij*_ = *H*_const_). The values in parentheses represent the fraction of snapshots with RMSD less than 1 Å.

### 4.3. Timestep dependency of complex stability

The dependency of complex stability on the integration timestep Δ*t* was examined to determine the optimal timestep for CGRig simulations. Five independent replicas of the CGRig simulations were conducted for each of the eleven protein dimer complexes using various timesteps. To assess stability, we calculated the average COM distances between the monomers and the Cα-RMSDs. Plots of these values against the timesteps revealed that the complex structures remained stable up to Δ*t* = 0.5 ps in all cases, and even at Δ*t* = 1 ps for seven of the eleven complexes (Figure 4). These results suggest that CGRig simulations can be safely performed with Δ*t* = 0.5 ps, while Δ*t* = 1 ps may be feasible depending on the system. Given that the optimal timestep varies across different protein systems, we recommend performing this stability test following NELVEX parameterization.

**Figure 4.**
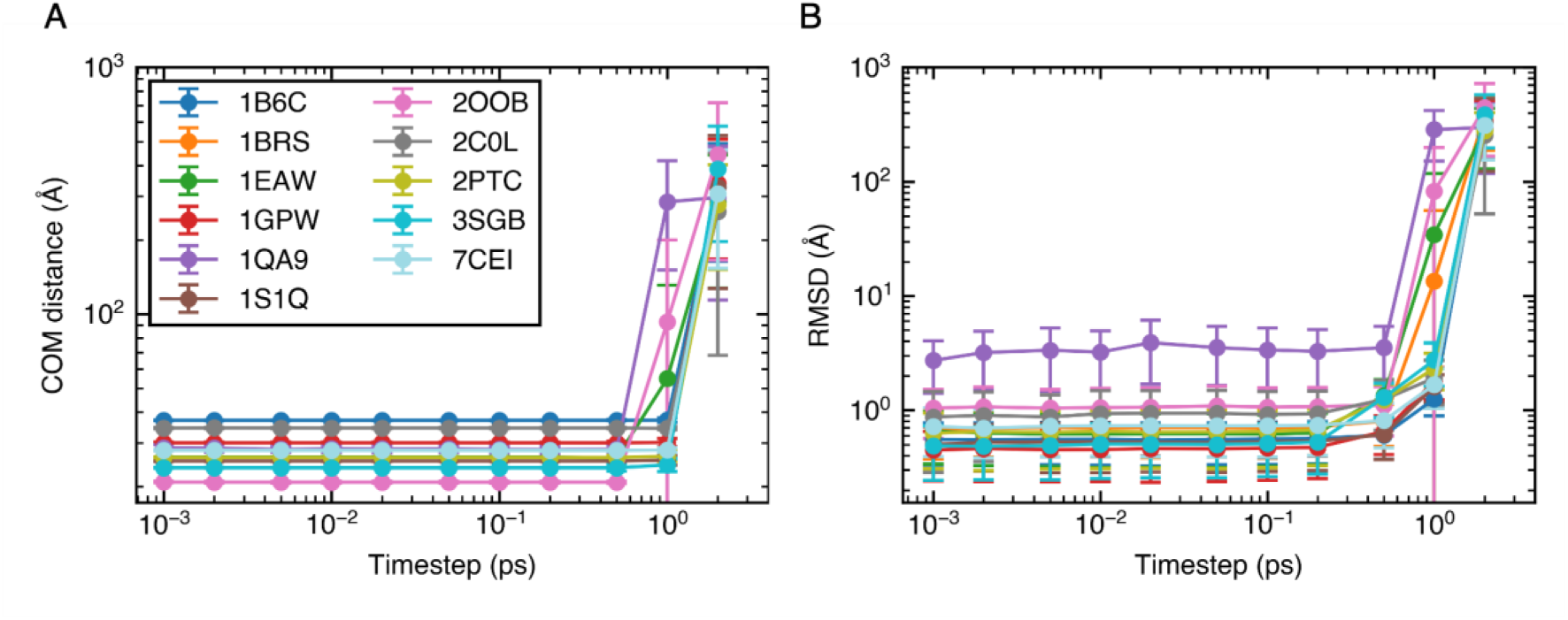
Dependence of structural stability on the integration timestep in CGRig simulations of eleven protein dimer complexes. (A) Average COM distances between the two protein subunits. (B) Average Cα-RMSD from the reference dimer structure. The RMSD calculation follows the same procedure as in Figure 2. Each data point and error bar represent the mean and standard deviation obtained from five independent simulations.

### 4.4. Protein association simulations

To evaluate whether CGRig can reproduce the association behavior from an unbound state, we performed simulations for the barnase–barstar system starting from a dissociated configuration. The two proteins were initially placed with a minimum distance of 35 Å. We conducted 50 independent replicas to observe the spontaneous formation of the complex and to ensure statistical reliability. Association events were observed following an initial diffusion period (Movie S1). To characterize the association process, we extracted an encounter ensemble by selecting snapshots in which the COM distance between the two proteins was less than 30 Å. These snapshots were then categorized into structural clusters. More than 60% of the snapshots in the encounter ensemble were classified into a single dominant cluster (Figure 5A). The representative structure of this cluster was remarkably similar to the reference complex structure, with an RMSD of less than 1 Å (Figure 5B). These results highlight the capability of CGRig to accurately recover the native binding mode, even when starting from a fully dissociated state.

**Figure 5.**
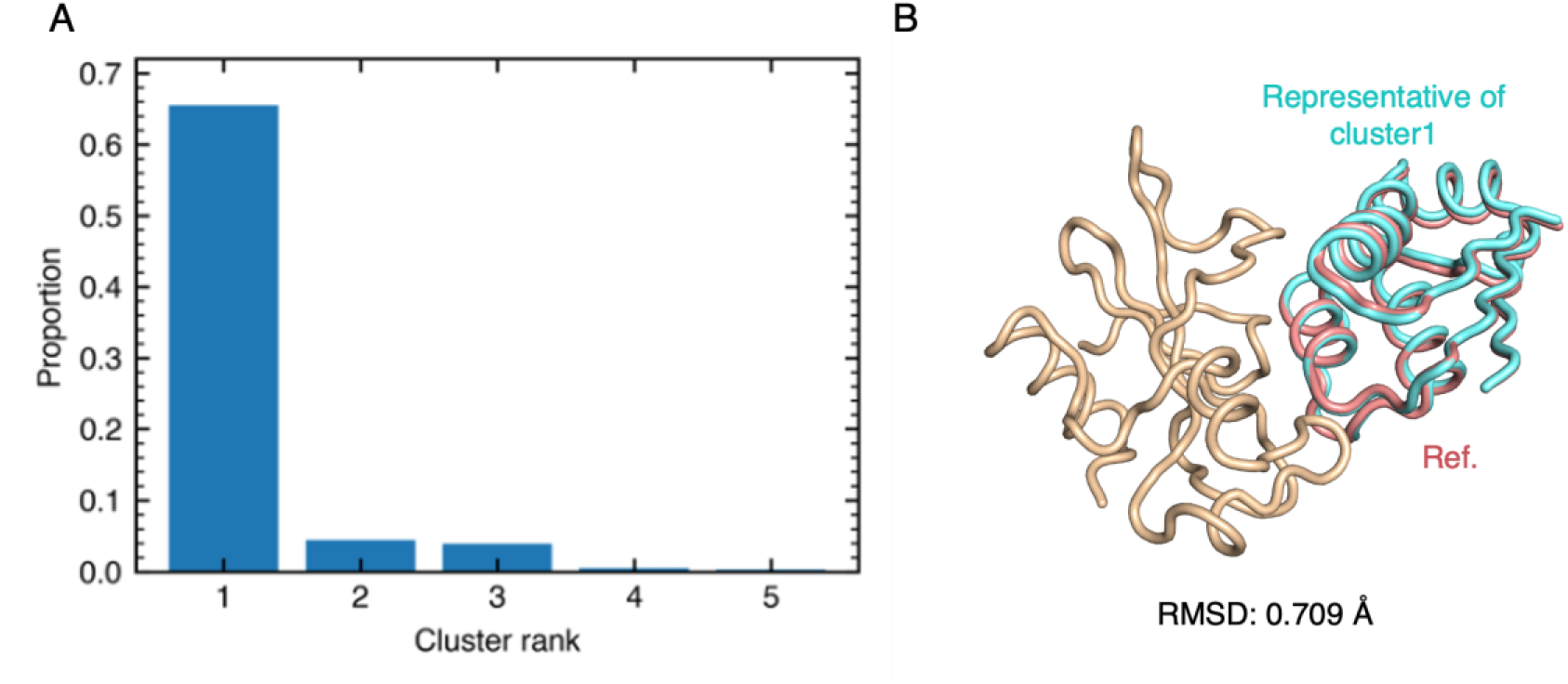
Protein association simulation results. (A) Fraction of snapshots assigned to the top five clusters. (B) Comparison between the representative structure of Cluster 1 and the reference structure. The barnase subunits (brown) are superimposed. The barstar subunit of the Cluster 1 representative structure is shown in red, while that of the reference structure is shown in blue.

We next estimated the association rate constant (*k*_on_ ) from the 50 trajectories. Following previous studies^70,71^, the two proteins were considered associated when native contacts were formed at two or more distinct sites. The calculated *k*_on_of (1.56 ± 0.28) × 10^9^ M^−1^s^−1^ is in excellent agreement with the value *k*_1_ = (1.8 ± 0.2) × 10^9^ M^−1^s^−1^ reported in a previous AA MD study^72^, which corresponds to the rate constant for encounter-complex formation (Table 2). Considering that the rapid association of barnase–barstar is primarily driven by electrostatic interactions^72^, these results demonstrate that CGRig accurately describes both the diffusion dynamics and the long-range electrostatics incorporated in the NELVEX potential. However, the calculated *k*_on_value is approximately five times larger than the experimental *k*_on_of (2.86 ± 0.67) × 10^8^ M^−1^s^−1 73^. This discrepancy is likely attributed to the omission of dehydration and induced-fit processes in CGRig simulations, which typically occur during the transition from an encounter complex to a tight complex.

**Table 2.** Comparison of calculated and experimental association rate constants for the barnase–barstar complex.

| Model | Rate constant<sup>a</sup> | Value<sup>b</sup> |
| --- | --- | --- |
| CGRig | $k_{\text{on}} (\text{M}^{-1}\text{s}^{-1})$ | $(1.56 \pm 0.28) \times 10^9$ |
| AA MD<sup>a</sup> | $k_1 (\text{M}^{-1}\text{s}^{-1})$ | $(1.8 \pm 0.2) \times 10^9$ |
| | $k_2 (\text{s}^{-1})$ | $(2.7 \pm 0.5) \times 10^{10}$ |
| | $k_{\text{on}} (\text{M}^{-1}\text{s}^{-1})$ | $(2.3 \pm 1.0) \times 10^8$ |
| Experiment | $k_{\text{on}} (\text{M}^{-1}\text{s}^{-1})$ | $(2.86 \pm 0.67) \times 10^8$ |

<sup>a</sup> $k_1$ , $k_2$ , and $k_{\text{on}}$ denote the rate constants of the formation of the encounter complex from the unbound state, the transition from encounter complex to bound state, and the overall association from the unbound to the bound state.
<sup>b</sup>Values are presented as means with 95% confidence intervals.

### 4.5. Tubulin self-assembly simulations

CGRig was motivated by the need to access long-time, large-scale protein self-assembly while accounting for protein-shape anisotropy. A representative target is the self-assembly of α/β tubulin dimers into microtubules^74^. In particular, the mechanism of the nucleation—the initial process of microtubule formation—is difficult to elucidate through experiments alone. Furthermore, performing AA MD simulations of large-scale protein self-assembly remains challenging due to high computational costs. Because tubulin oligomerization involves distinct longitudinal and lateral interfaces (see Figure S1), spherical approximations are ill-suited for describing these specific, anisotropic interactions. Therefore, we tested whether CGRig can reproduce the oligomerization process of α/β tubulin dimers.

We performed a CGRig simulation for a system composed of 16 tubulin dimers (32 subunits), which were initially placed on a lattice (Figure 6). The tubulin dimers rapidly associated to form tetramers, followed by further oligomerization, ultimately producing a predominantly longitudinal oligomer containing nine tubulin dimers. The α- and β-tubulin subunits remained associated as dimers throughout the simulation. This preference for longitudinal association is consistent with previous computational studies^75,76^ and an experimental model^77^. These results demonstrate the potential of CGRig for probing early-stage nucleation mechanisms over extended spatiotemporal scales.

**Figure 6.**
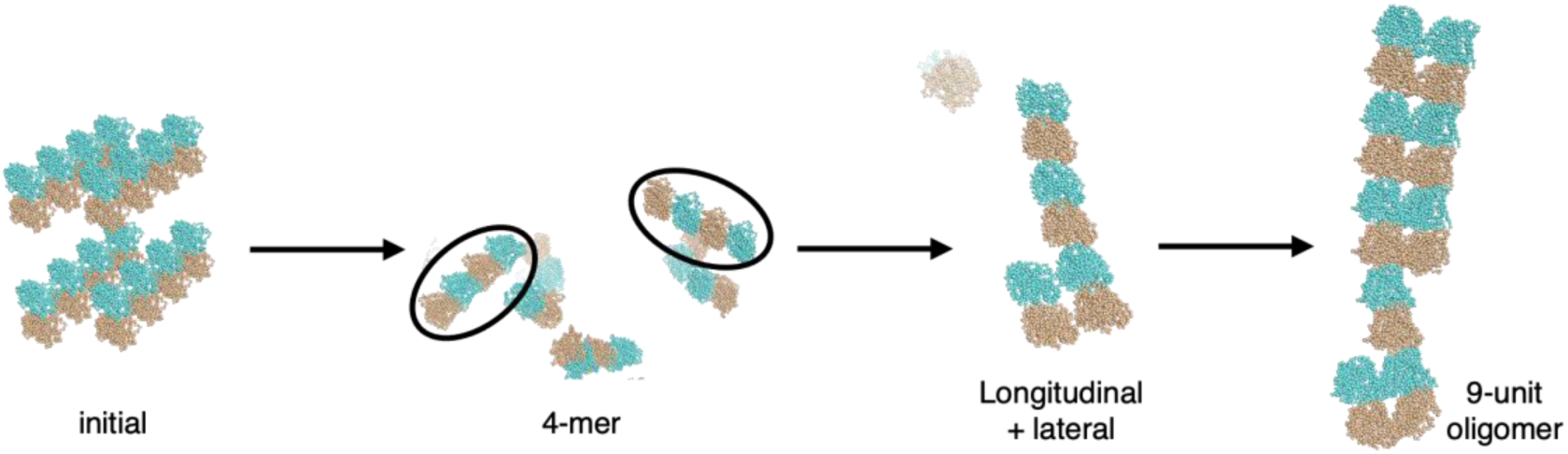
Snapshots of the tubulin self-assembly process. The images show the sequence from the initial state to the formation of a 9-unit oligomer via tetramer and small longitudinal intermediates. α- and β-tubulin subunits are colored brown and blue, respectively.

### 4.6. Performance and scaling

Finally, we evaluated computational efficiency as a function of system size (number of tubulin subunits) and compared the performance between CPU-only and GPU-enabled implementations (Figure 7). As expected, GPU acceleration proved significantly more effective as the system size increased. For a system comprising 1,024 tubulin subunits, the GPU-enabled implementation achieved 205.894 steps/s. With Δ*t* = 1 ps, this performance corresponds to 17.8 *μ*s/day. This implies that a 1-ms simulation of a system with over 1,000 subunits can be completed in approximately two months using a single GPU.

**Figure 7.**
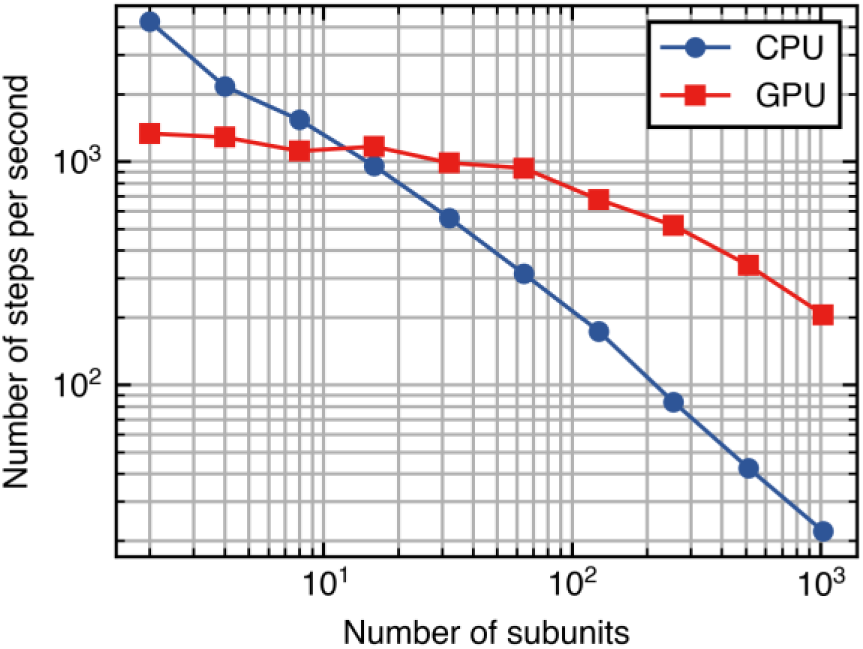
Performance benchmark results. Simulation throughput (steps per second) as a function of the number of subunits for CPU-only and GPU-enabled implementations.

## 5. CONCLUSIONS

In this study, we developed the CGRig framework, which treats protein molecules as rigid bodies with embedded interaction sites. The dynamics of these rigid bodies are described by overdamped Langevin equations incorporating full (6 × 6) friction matrices derived from the all-atom coordinates. This approach explicitly accounts for the coupling between translational and rotational motions. While the CGRig framework can embed interaction sites at an arbitrary coarse-graining level, we implemented a residue-level interaction potential termed NELVEX (Native contact, Electrostatics, and Volume Exclusion). This potential combines a Go̅-like native contact potential, optimized via force-matching against reference AA MD simulations, with Debye–Hückel electrostatics and volume exclusion. With the NELVEX potential, protein complex structures were stably maintained throughout the CGRig simulations. Furthermore, protein association simulations demonstrated that CGRig successfully recovers the native complex structure even when starting from a fully dissociated state. The model also yielded a reasonable association rate, confirming the correct description of both diffusion dynamics and long-range electrostatics. The CGRig simulations are stably performed with a timestep (Δ*t*) of 0.5 ps, and Δ*t* = 1.0 ps is also feasible for certain proteins. Specifically, the model’s applicability was demonstrated through a tubulin self-assembly simulation, where the oligomerization of dimers was reproduced consistently with previous studies. In terms of computational efficiency, the GPU-enabled implementation of CGRig achieved a throughput of 17.8 μs/day for a system of 1,024 subunits. Consequently, this framework allows for significant spatiotemporal expansion while retaining the structural specificity necessary to describe complex protein assembly process.

While the current framework has proven highly effective, several limitations remain to be addressed in future work. For instance, the omission of desolvation effects and induced-fit processes during protein association resulted in association rates larger than those observed experimentally. To address this issue, incorporating Monte Carlo methods to stochastically adjust intermolecular interactions may provide a viable solution. Furthermore, considering that many proteins contain functionally significant flexible regions or intrinsically disordered regions (IDRs), the current rigid-body framework has inherent limitations. To address this, a multi-resolution approach^78,79^ that integrates rigid bodies for folded domains with flexible particle-based representations for disordered segments should be pursued in future development of CGRig. Despite these challenges, the CGRig framework established in this study provides a robust foundation for simulating large-scale protein assemblies, offering a practical balance between computational efficiency and structural detail.

## ASSOCIATED CONTENT

### Supporting Information

Additional figure and analysis of the CGRig simulations and AA MD simulations (PDF)

Trajectory of barnase–barstar association simulation (MP4)

Trajectory of tubulin self-assembly simulation (MP4)

## AUTHOR INFORMATION

### Author Contributions

Y.T. and T.T. designed the research. Y.T. implemented the source code and performed the simulations. All authors analyzed the results and wrote the manuscript.

### Notes

The authors declare no competing financial interest.

## ACKNOWLEDGMENT

Y. T. acknowledges JST SPRING, Grant Number JPMJSP2108, and Initiative on Recommendation Program for Young Researchers and Woman Researchers, Information Technology Center, The University of Tokyo. This research was partially supported by Research Support Project for Life Science and Drug Discovery (Basis for Supporting Innovative Drug Discovery and Life Science Research (BINDS)) from AMED under Grant Number JP25ama121027. The authors used Gemini (Google) for linguistic refinement of the manuscript.

