## Supplementary material for "CGRig: a rigid-body protein model with residue-level interaction sites for long-time and large-scale protein assembly simulation": Figure S1, Figure S2, Figure S3, Figure S4

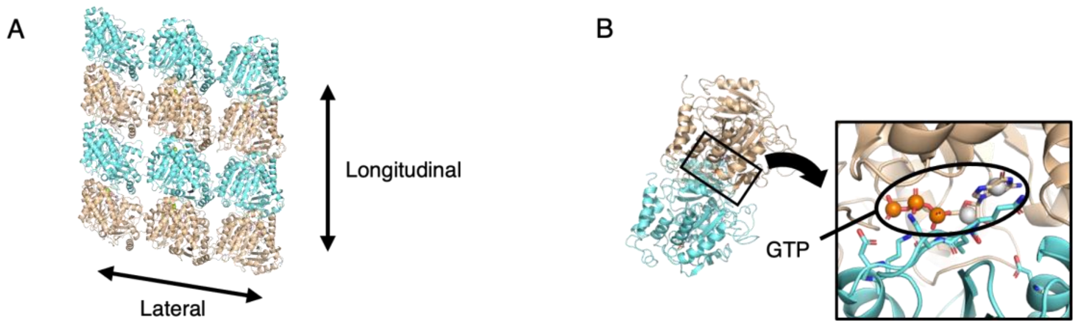


**Figure S1.** (A) Experimental structure of a tubulin 16-mer complex (PDBID: 6EVW). Horizontal and vertical arrows indicate lateral and longitudinal intermolecular interactions, respectively. (B) Modeling scheme for GTP. The close-up view shows GTP represented by five interaction sites located at the centers-of-mass (COMs) of the α-, β-, and γ-phosphate groups, as well as the base and sugar moieties.

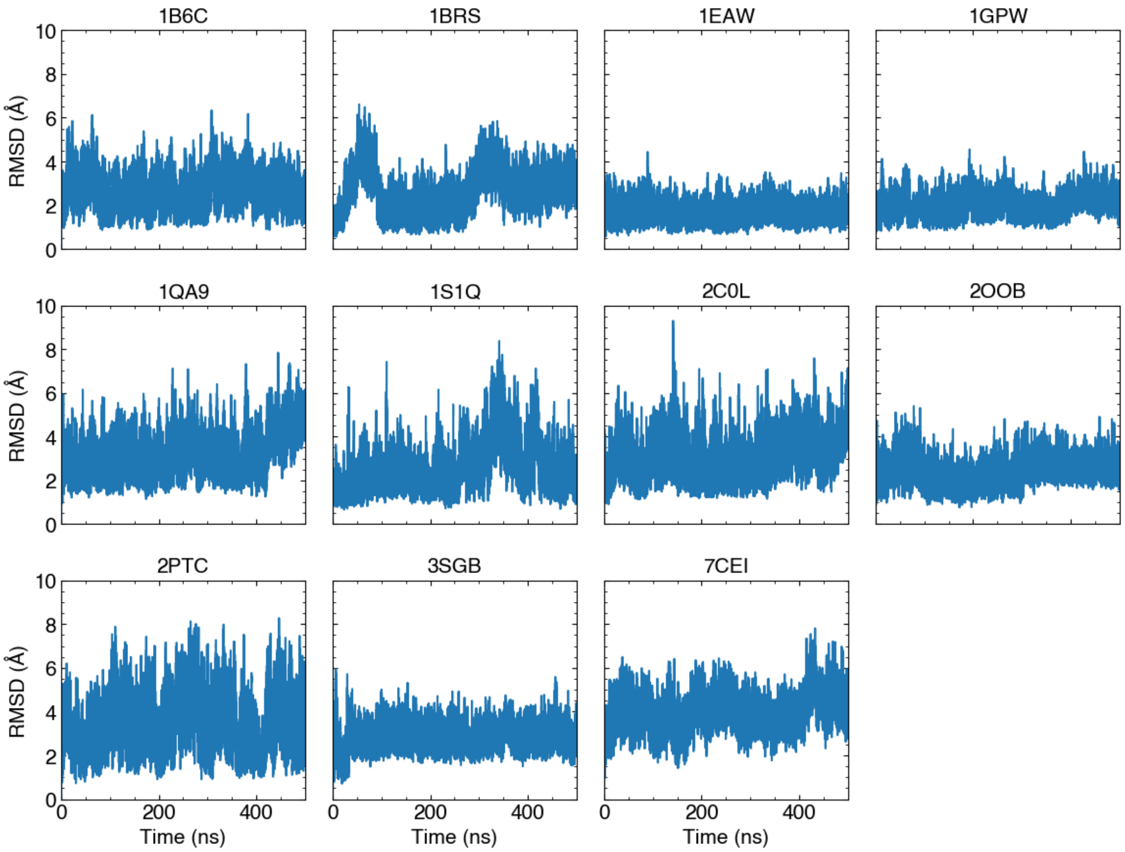


**Figure S2.** Time evolutions of Cα-RMSDs calculated from 500-ns AA MD simulations of eleven protein dimer complexes. The RMSD calculation follows the same procedure as in Figure 2. The PDB ID for each complex is shown above the corresponding panel.

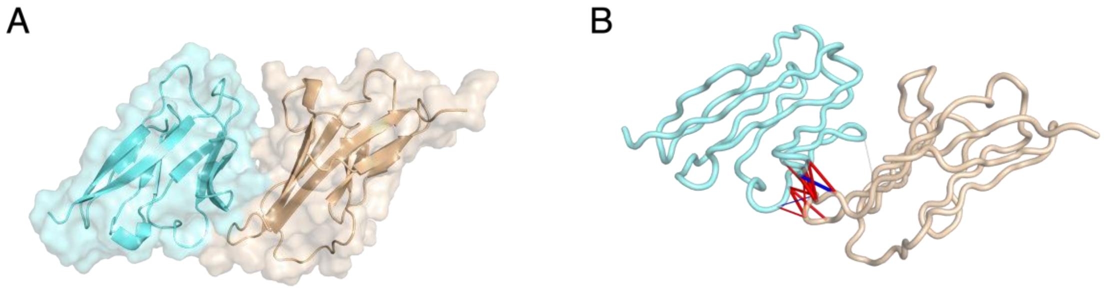


**Figure S3.** Structural details of 1QA9. (A) Backbone structures shown in cartoon representation with molecular surfaces. The larger (longer amino acid sequence) and smaller (shorter sequence) subunits are colored brown and blue, respectively. (B): Native contact pairs shown with the backbone structures. Red and blue lines represent $H_{ij} < 0$ and $H_{ij} > 0$, respectively, where the line thickness indicates the strength of the interaction.

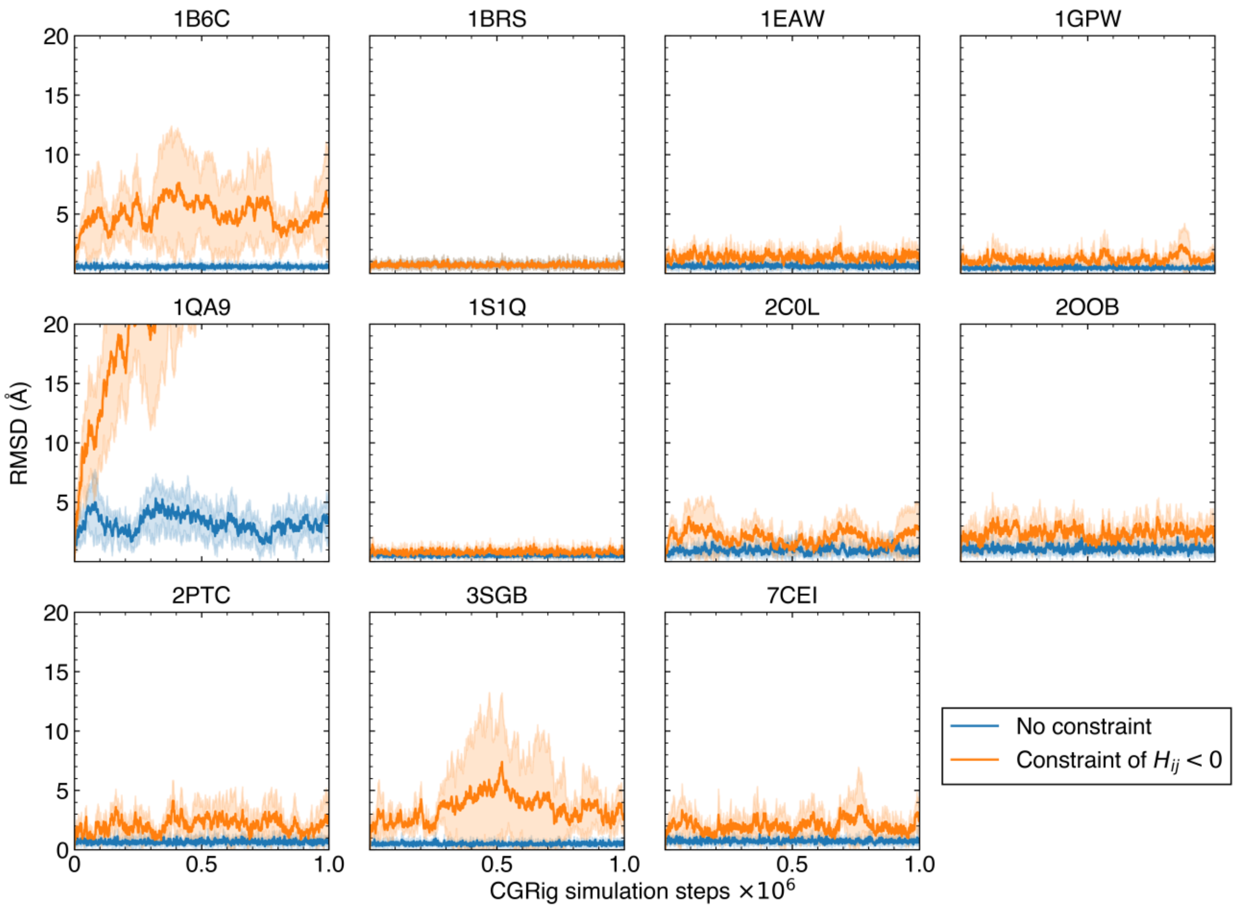


**Figure S4.** Time evolutions of Cα-RMSDs calculated from CGRig simulations, comparing the results using the NELVEX potential without constraints (blue) and with constraints of $H_{ij} < 0$ (orange). The RMSD calculation follows the same procedure as in Figure 2. The PDB ID for each complex is shown above the corresponding panel. Solid lines and shaded areas represent the mean values and standard deviations of five independent runs, respectively.
